# Identification of low micromolar SARS-CoV-2 M^pro^ inhibitors from hits identified by *in silico* screens

**DOI:** 10.1101/2020.12.03.409441

**Authors:** Giacomo G. Rossetti, Marianna Ossorio, Samia Barriot, Laurence Tropia, Vasilis S. Dionellis, Christoph Gorgulla, Haribabu Arthanari, Peter Mohr, Remo Gamboni, Thanos D. Halazonetis

## Abstract

M^pro^, also known as 3CL^pro^, is the main protease of the SARS-CoV-2 coronavirus and, as such, is essential for the viral life cycle. Two studies have each screened and ranked *in silico* more than one billion chemical compounds in an effort to identify putative inhibitors of M^pro^. More than five hundred of the seven thousand top-ranking hits were synthesized by an external supplier and examined with respect to their activity in two biochemical assays: a protease activity assay and a thermal shift assay. Two clusters of chemical compounds with M^pro^ inhibitory activity were identified. An additional five hundred molecules, analogues of the compounds in the two clusters described above, were also synthesized and characterized *in vitro*. The study of the analogues revealed that the compounds of the first cluster acted by denaturing M^pro^ and might denature other proteins as well. In contrast, the compounds of the second cluster targeted M^pro^ with much greater specificity and enhanced its melting temperature, consistent with the formation of stable M^pro^-inhibitor complexes. The most active compounds of the second cluster exhibited IC_50_ values between 4 and 7 μM and their chemical structure suggests that they could serve as leads for the development of potent M^pro^ inhibitors.

## INTRODUCTION

At the end of December 2019, an outbreak of pneumonia of initially unknown etiology was reported by the Chinese health authorities. A novel coronavirus was isolated from human airway epithelial cells and identified as the cause of a disease, now referred to as COVID-19, that produces a wide range of symptoms, including fever, cough, shortness of breath and loss of smell and taste as the most common ones [1]. On 11 March 2020, the World Health Organization declared the outbreak as a pandemic. As of December 1, 2020, the coronavirus pandemic has already infected more than 64 million people and caused more than 1.5 million deaths (www.worldometers.info/coronavirus/).

The identified novel coronavirus, first named 2019-nCoV and then SARS-CoV-2, is very similar to SARS-CoV (aka SARS-CoV-1) [2]. Coronaviruses are enveloped, positive-sense, single-stranded RNA viruses; the size of their genome is about 30,000 nucleotides and encodes at least six open reading frames (ORFs) [3]. Two large ORFs, ORF1a and ORF1b give rise to polyproteins, pp1a and pp1ab, respectively. A main protease (M^pro^, also called 3C-like protease) and a papain-like protease (PL^pro^) process the polyproteins pp1a and pp1ab into 16 nonstructural proteins, which are important for synthesis of viral RNA and structural proteins (envelope, membrane, spike and nucleocapsid proteins). The three-dimensional structure of SARS-CoV-2 M^pro^ has been described by several groups [4–9], revealing that the N-terminus of M^pro^ contains two domains (domain I, residues 10-99; and domain II, residues 100-184) that adopt a chymotrypsin-like fold. The active site is located at the cleft formed between these domains and contains a Cys145-His41 catalytic dyad. A total of five protein segments comprising residues 25-27 and 44-50 from domain I and residues 140-143, 165-168 and 188-190 from domain II form the walls of the active site.

The M^pro^ protease is essential for the viral life cycle and is also among the most highly conserved proteins in the coronavirus family. For example, the M^pro^ proteases of SARS-CoV-1 and SARS-CoV-2 are 96% identical at the amino acid sequence level and their active sites are 100% identical [4]. For the above reasons, M^pro^ is considered to be a good target for the development of novel treatments for COVID-19. Effective inhibitors of SARS-CoV-2 M^pro^ could impact the course of COVID-19, but, more importantly, they might prevent possible future pandemics caused by other Betacoronaviruses.

The conservation of M^pro^ extends beyond the coronavirus family. Thus, GC376, one of the most potent inhibitors of SARS-CoV-2 M^pro^, was originally developed as an inhibitor of the 3CL^pro^ of Norwalk virus, a member of the calicivirus family [10]. GC376 and the related compounds GC373 and GC375 are dipeptidyl compounds with different warheads [10–12]. The electrophilic warheads allow these inhibitors to form covalent bonds with the thiol group of the active site cysteine. GC376 inhibits the main proteases of picornaviruses, caliciviruses and coronaviruses [10] and is being developed for the treatment of cats infected with feline coronavirus [13]. GC376 also inhibits the M^pro^ of SARS-CoV-2 with an IC_50_ of 30 ± 8 nM [14,15].

Another potent inhibitor of SARS-CoV-2 M^pro^, PF-00835231, was developed originally as an inhibitor of the M^pro^ of SARS-CoV-1 [16]. PF-00835231 is structurally related to GC376, but features a different N-capping group and uses a hydroxy-methyl-ketone as warhead to form a covalent bond with the active site cysteine; it has an impressive *in vitro* IC_50_ of 0.27 ± 0.1 nM [16]. However, efforts to develop this compound for treatment of COVID-19 in humans suggest that a continuous intravenous infusion of a prodrug would be needed to achieve effective doses in the plasma of human patients [17].

Additional efforts to develop SARS-CoV-2 M^pro^ inhibitors include screening by X-ray crystallography of a library of very small chemical compounds (referred to as fragments) to identify compounds that bind M^pro^ [18,19]. These hits can be used as starting points to develop M^pro^ inhibitors.

Finally, many groups have used the reported crystal structure of M^pro^ to identify inhibitors by *in silico* screening. Two of these studies screened more than one billion compounds [20,21]. As our group participated in one of these endeavors, we followed upon this work by examining whether some of the top hits could actually inhibit the protease activity of SARS-CoV-2 M^pro^ *in vitro*. We describe here our experience and the identification of three structurally related inhibitors that inhibit SARS-CoV-2 M^pro^ *in vitro* with IC_50_ values between 4.2 and 7.4 μM.

## MATERIALS AND METHODS

### Protein expression and purification of SARS-CoV-2 M^pro^

The M^pro^ construct [4] provided by Rolf Hilgenfeld was transformed into *E. coli* strain BL21-Gold (DE3) (Agilent). Transformed clones were picked to prepare pre-starter cultures in 2 mL YT medium with ampicillin (100 μg/ml), at 37°C for 8 h. The pre-starter culture was then inoculated into fresh 120 mL YT medium with ampicillin (100 μg/ml) and incubated at 37°C overnight. The next day, the starter culture was inoculated into 1,600 mL YT medium with ampicillin (100 μg/ml) and incubated at 37°C until OD_600_ reached a value between 0.6 and 0.8. 1 mM isopropyl-D-thiogalactoside (IPTG) was then added to induce the overexpression of M^pro^ at 30°C for 5 h. The bacteria were harvested by centrifugation at 8260 x g, 4°C for 15 minutes, resuspended in Binding Buffer (25 mM BTP pH6.8; 300 mM NaCl; 2 mM DTT; 1 mM EDTA; 3% DMSO) and then lysed using an Emulsiflex-C3 homogenizer (Avestin). The lysate was clarified by ultracentrifugation at 137,088 x g, 4°C for 1 h and loaded onto a HisTrap FF column (Cytiva) using an Äkta protein purification system (Cytiva). When all the supernatant containing M^pro^ had passed through the column, the column was washed with 80 mL binding buffer to remove non-specifically bound proteins and then M^pro^ was eluted using an imidazole gradient (0-500 mM) in Binding Buffer. The M^pro^ fractions were concentrated using 3 kDa Amicon Ultra Centrifugal Filters (Merck Millipore) and the M^pro^ protein was further purified by size exclusion chromatography using a HiLoad Superdex 200 column (Cytiva) attached to a SMART protein purification system (Pharmacia).

### Compounds

482 compounds, referred to as parent compounds, were selected from lists of high-ranking, based on *in silico* screens, putative M^pro^ inhibitors (Supplementary Tables 1-4). After evaluating the activity of these compounds *in vitro*, we selected an additional 578 compounds that were analogues of a few active parent compounds (Supplementary Tables 5-7). All the above compounds were purchased from Enamine, their purity was ≥ 90% and most of them were synthesized on customer’s demand. The compounds were dissolved in DMSO at a concentration of 2 mM and were stored at −20°C. GC376 was purchased from BPSBioscience.

### SARS-CoV-2 M^pro^ protease activity assay

M^pro^ protease assays were performed in duplicate in Falcon 384-well optilux flat bottom, TC-treated microplates (Corning) in a final volume of 10 μL. M^pro^ protease, at a final concentration of 100 nM, was preincubated for 20 min at room temperature (RT) with the compounds in assay buffer (5 mM HEPES pH 7.5, 0.1 mg/mL BSA, 0.01% Triton, 2 mM DTT) under gentle agitation. The FRET substrate, HiLyte-Fluor488-ESATLQSGLRKAK(QXL520)-NH2 (Eurogentec), was then added at a final concentration of 500 nM and incubated for 2 min at RT with gentle agitation prior to the start of fluorescence measurement. Compounds and FRET substrate were dispensed with an acoustic liquid dispenser (Gen5-Acoustic Transfer System; EDC Biosystems). The fluorescence intensity was measured kinetically for 7 cycles, every 10 min at 22°C, using a Spark 10M microplate reader (Tecan) and excitation and emission wavelengths of 485 and 528 nm, respectively.

### SARS-CoV-2 M^pro^ thermal shift binding assay

The binding of the compounds to M^pro^ was monitored by differential scanning fluorimetry (DSF). M^pro^ protease thermal shift assays were performed in duplicate in LightCycler 480 multiwell plates 96, white (Roche) in a final volume of 20 μL. M^pro^ protease, at a final concentration of 1 μM, was preincubated for 20 min at room temperature (RT) under gentle agitation with the compounds (final concentration: 20 μM) in assay buffer (10 mM HEPES pH 7.5; 150 mM NaCl). Protein melting was monitored using 1000X SYPRO Orange (Sigma) binding dye. Compounds and SYPRO Orange were dispensed with an acoustic liquid dispenser (Gen5-Acoustic Transfer System; EDC Biosystems). Fluorescence (excitation wavelength: 465 nm; emission wavelength: 580 nm) was measured over a temperature gradient ranging from 20 to 95 °C, with incremental steps of 0.05 °C/s and 11 acquisitions per °C. The melting curves and peaks were obtained using the melting temperature (Tm) calling analysis of the LightCycler 480 Software (release 1.5.1.62; Roche).

## RESULTS AND DISCUSSION

### Development of a FRET-based assay for SARS-CoV-2 M^pro^ activity

The plasmid encoding the SARS-CoV-2 M^pro^ protease with a C-terminal His-tag was kindly provided by Rolf Hilgenfeld [4]. M^pro^ was expressed in *E. coli* strain BL21-Gold (DE3) and the recombinant protein was purified in two steps by affinity and size exclusion chromatography (Fig. 1a).

**Fig. 1.**
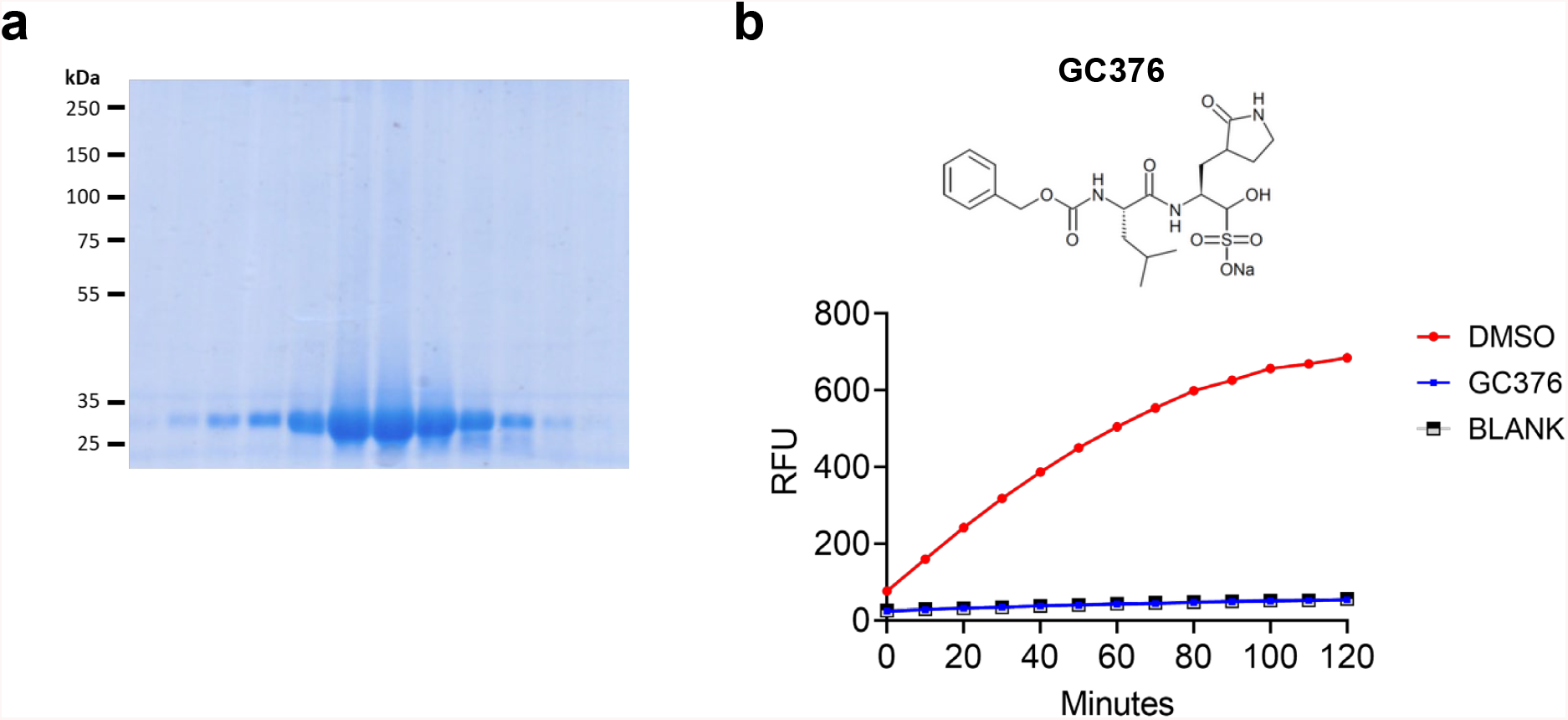
SARS-CoV-2 M^pro^ expression and protease activity. **a.** SDS-PAGE of fractions of His-tagged M^pro^ eluted by an imidazole gradient from a HisTrap FF column. The horizontal ticks indicate the positions of migration of the molecular weight markers. The purified M^pro^ protein migrates at about 30 kDa, consistent with its molecular weight. **b.** The protease activity of M^pro^ was monitored by a FRET-assay, as described in the text. In this experiment, fluorescence was measured every 10 min over a period of 120 min. GC376, a previously-described M^pro^ inhibitor, whose structure is shown, was examined at a final concentration of 40 μM. RFU, relative fluorescence units.

The protease activity of purified M^pro^ was studied using a FRET-based assay suitable for high throughput analysis. The FRET substrate peptide contains at its N-terminus a dye (HiLyte-Fluor488), whose fluorescence is quenched by a C-terminal quencher (QXL520). Cleavage of the peptide by M^pro^ leads to an increase in fluorescence intensity. The assay was carried out in 384-well plates in a reaction volume of 10 μL with the protease and fluorogenic substrate at final optimized concentrations of 100 nM and 500 nM, respectively. The fluorescence intensity was measured kinetically every 10 min in a microtiter plate-reading fluorimeter in the presence or absence of the M^pro^ inhibitor GC376 [10–12]. In the absence of the inhibitor, the fluorescence intensity increased linearly during the first 60 min of the reaction, whereas in the presence of 40 μM GC376 no increase in fluorescence intensity was observed (Fig. 1b). The substrate and inhibitor were dispensed using an acoustic liquid dispenser, which allowed compounds to be dispensed directly from the stock solution plates.

### Validation of putative SARS-CoV-2 M^pro^ inhibitors identified by *in silico* screens

We described recently two *in silico* screens of one billion compounds using M^pro^ as a target [20]. The first screen, hereafter referred to as screen 1A, used the three-dimensional coordinates of M^pro^ described by Jin et al [5] (pdb id: 6lu7). The second screen, hereafter referred to as screen 1B, used the coordinates of M^pro^ described by Dai et al [3] (pdb id: 6m0k), except that different rotamers were used for residues S46, M49 and C145, in the hope of expanding the active site and capturing a larger repertoire of chemical compounds [20]. Comparison of the top 1,000 hits identified by screens 1A and 1B, revealed an overlap of only 12 compounds.

From screen 1A, 3,808 top-ranking hits were evaluated and 195 compounds thereof were chosen to be synthesized (Table S1). Main selection criteria included drug-likeness and chemical diversity. From screen 1B, 3,851 top hits were evaluated and 226 compounds thereof were selected for chemical synthesis (Table S2). In addition, guided by the results of a crystallographic fragment screen [19] that showed a fragment containing a nitrile group deep in the active site of M^pro^ (pdb id: 5r82), we identified all the nitrile-containing compounds among the top 20,000 hits of screens 1A and 1B. This list included 253 compounds, 45 of which were synthesized (Table S3). Finally, we also ordered synthesis of 12 of the top 15 hits from screen 1A and of 8 of the top 15 hits from an *in silico* screen of 1.3 billion compounds conducted by Ton et al [21], hereafter referred to as screen 2 (Table S4).

All the above compounds, 486 in total, were assayed at a final compound concentration of 40 μM for their ability to inhibit the protease activity of M^pro^. From the 207 compounds selected from the hits of screen 1A (Table S1 and first 12 compounds of Table S4), only weak inhibitors of M^pro^ were identified and none of them were pursued further. From the 226 compounds selected from the hits of screen 1B (Table S2), one compound, Z1037455358, was particularly active (Fig. 2a,c); whereas from the 45 nitrile-containing compounds selected from the hits of screens 1A and 1B (Table S3), the structurally-related compounds Z637352244 and Z637352642 (both from screen 1A) appeared promising (Fig. 2a,c). Finally, from the 8 compounds selected from the hits of screen 2 (bottom 8 compounds of Table S4), two structurally-related compounds, ZINC000636416501 and ZINC000373659060, were weakly active (Fig. 2b,c). We decided to investigate further the compounds cited above by having synthesized or ordering, when commercially available, analogous compounds based on similarity and substructure searches.

**Fig. 2.**
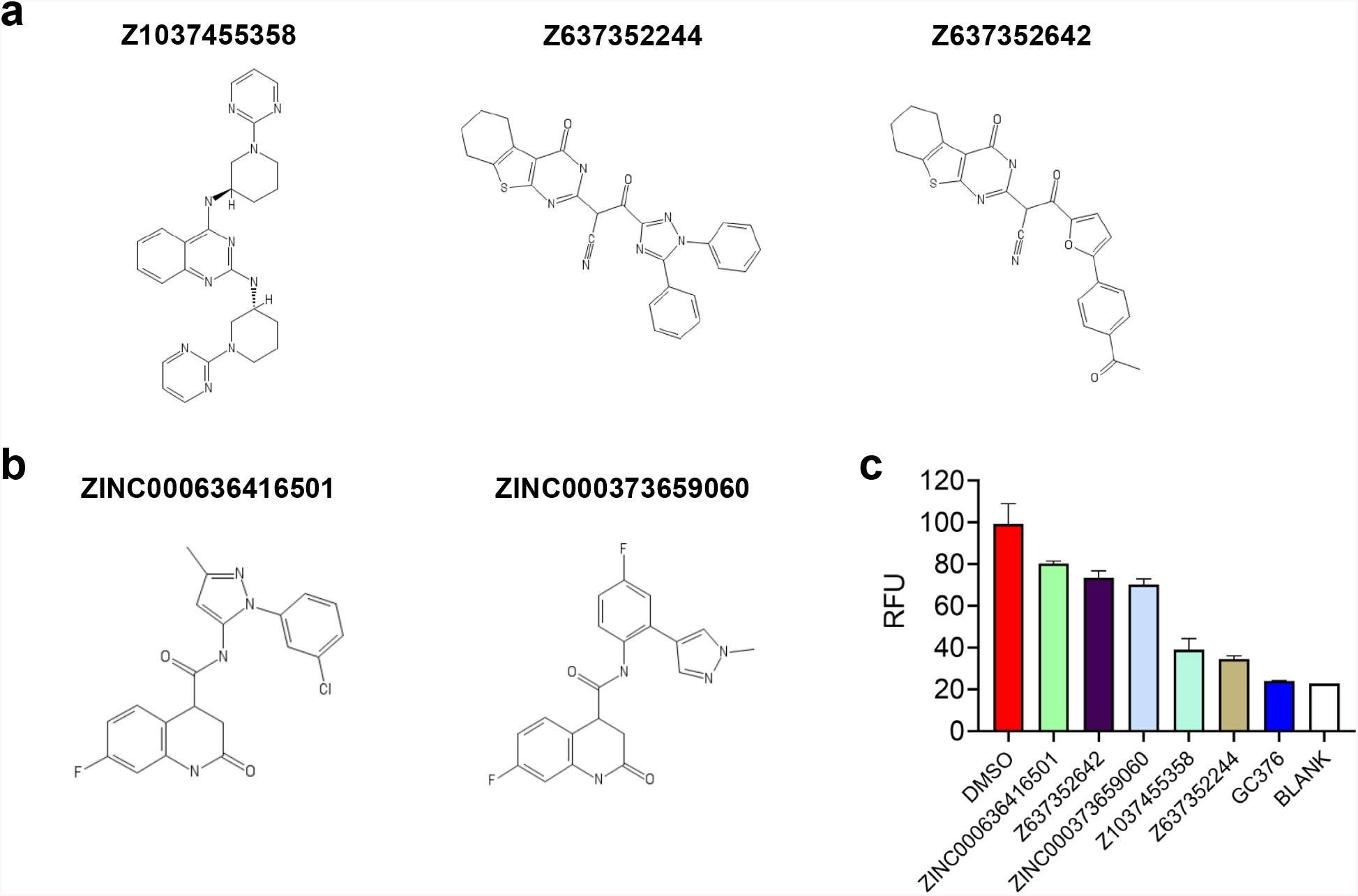
Identification of compounds that inhibit the protease activity of SARS-CoV-2 M^pro^. **a.** Chemical structures of the most active compounds from screens 1A and 1B. **b.** Chemical structures of the most active compounds from screen 2. **c.** Graph showing the inhibitory activity of the compounds tested at a final concentration of 40 μM. After addition of the FRET substrate, fluorescence was acquired at 10 min intervals over 60 min. For each compound, the increase in fluorescence intensity was normalized to the DMSO control. GC376 serves as a positive control. RFU, relative fluorescence units.

### Characterization of a cluster of nitriles and a diamino-quinazoline singleton

We first focused our effort on the two structurally-related compounds Z637352244 and Z637352642 (Fig. 2c). These two compounds were identified by screen 1A and have in common a nitrile, a functional group which is also present in a fragment that was found to bind M^pro^ by crystallographic screening [18]. To identify more potent compounds, we ordered 301 analogues, most of which were chemically synthesized for this project (Table S5). As expected, several of the analogues inhibited M^pro^, when tested at a final concentration of 40 μM (Fig. 3a). We determined the IC_50_ concentrations of the original compounds and of the five most promising analogues. The parent compounds Z637352244 and Z637352642 exhibited IC_50_ values of 16 and 96 μM, respectively (Fig. 3b,c). Three analogues, Z56785964, Z637450230 and Z56786187, showed IC_50_ values between 13-17 μM, whereas two analogues, Z2239054061 and Z637352638, had IC_50_ values of 6.7 and 7.5 μM, respectively (Fig. 3b,d). We will refer to this family of compounds as the cluster of nitriles.

**Fig. 3.**
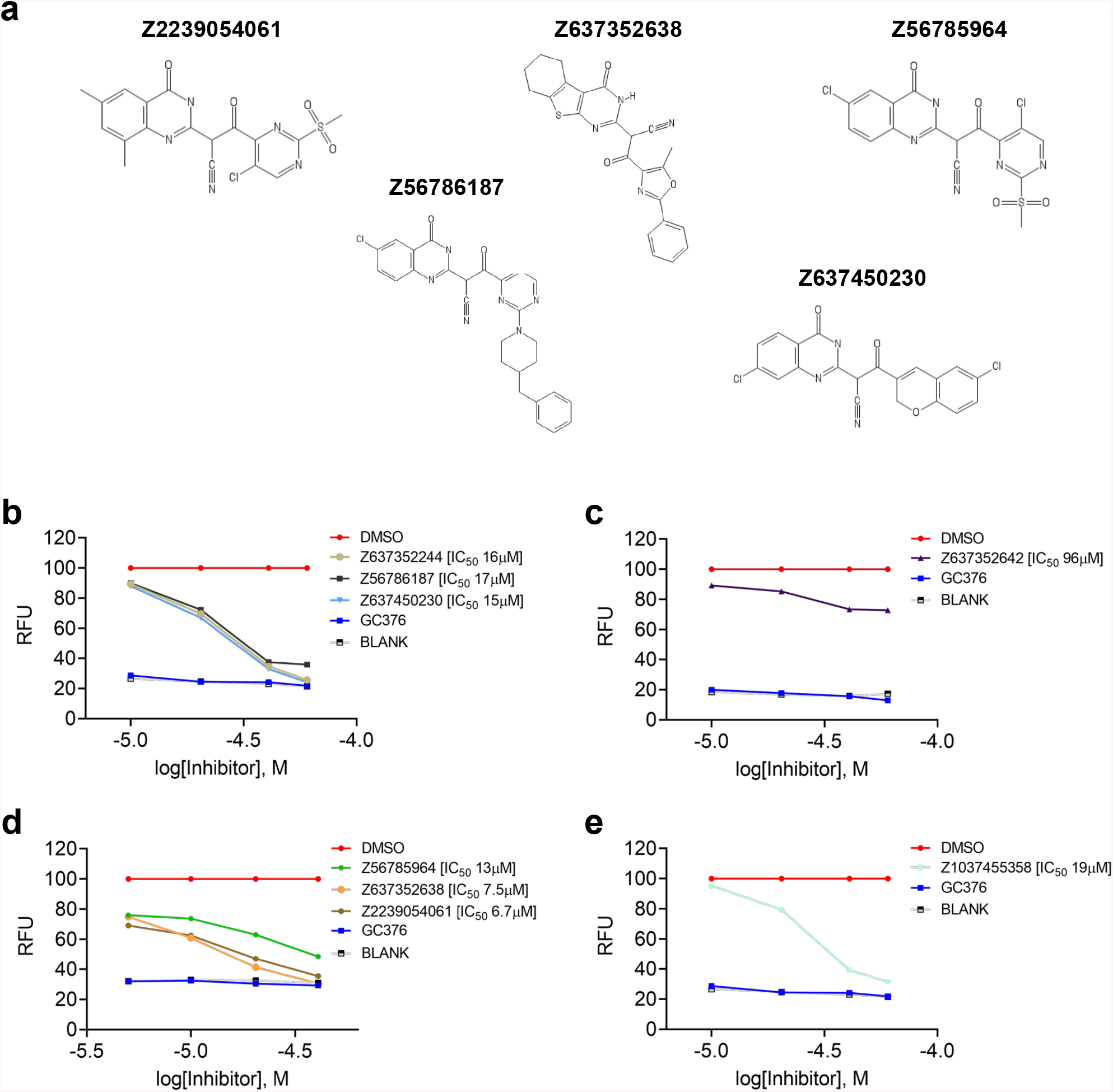
Inhibition of M^pro^ protease activity by compounds containing nitrile or diamino-quinazoline groups. **a.** Chemical structures of the most active analogues from the cluster of nitriles. **b-d.** Dose-response curves for compounds Z637352244, Z637352642 and their analogues examined at 10, 20, 40 and 60 μM final compound concentrations (**b,c**) and 5, 10, 20 and 40 μM final compound concentrations (**d**). The protease assay was performed, as described in Fig. 2c. RFU, relative fluorescence units. **e.** Dose-response curve for the diamino-quinazoline compound Z1037455358 examined at 10, 20, 40 and 60 μM final compound concentrations.

We next focused our efforts on compound Z1037455358, which was identified by screen 1B and which contained a diamino-quinazoline core. The IC_50_ of this compound was 19 μM (Fig. 3e). We obtained 108 analogues of this compound (Table S6), but none of them were more potent than the parent compound and, therefore, none of them were further pursued. However, we retained the parent compound for further analysis and we will refer to it as the diamino-quinazoline singleton.

To continue validation of the nitrile cluster and the diamino-quinazoline singleton, we examined their effect in a thermal shift assay (TSA). Briefly, M^pro^ protease, at a final concentration of 1 μM, was incubated for 20 min with the inhibitors at a final concentration of 20 μM; the melting temperature of M^pro^ was then determined. Compounds that bind to M^pro^ should enhance its melting temperature [22]. Indeed, GC376 increased the melting temperature of M^pro^ by 19°C (Fig. 4a). Surprisingly, the parent nitrile-containing compounds Z637352244 and Z637352642 decreased the melting temperature of M^pro^ by 2-3°C (Fig. 4a). Three of their analogues, Z637352638, Z2239054061 and Z56785964, decreased the melting temperature of M^pro^ even more, by 8-11°C. Of the remaining two analogues, Z56786187 decreased the melting temperature of M^pro^ by 4°C, whereas Z637450230 had no effect on the melting temperature (Fig. 4a). Compound Z1037455358, the diamino-quinazoline singleton, also led to a modest decrease in the melting temperature of M^pro^ (Fig. 4b).

**Fig. 4.**
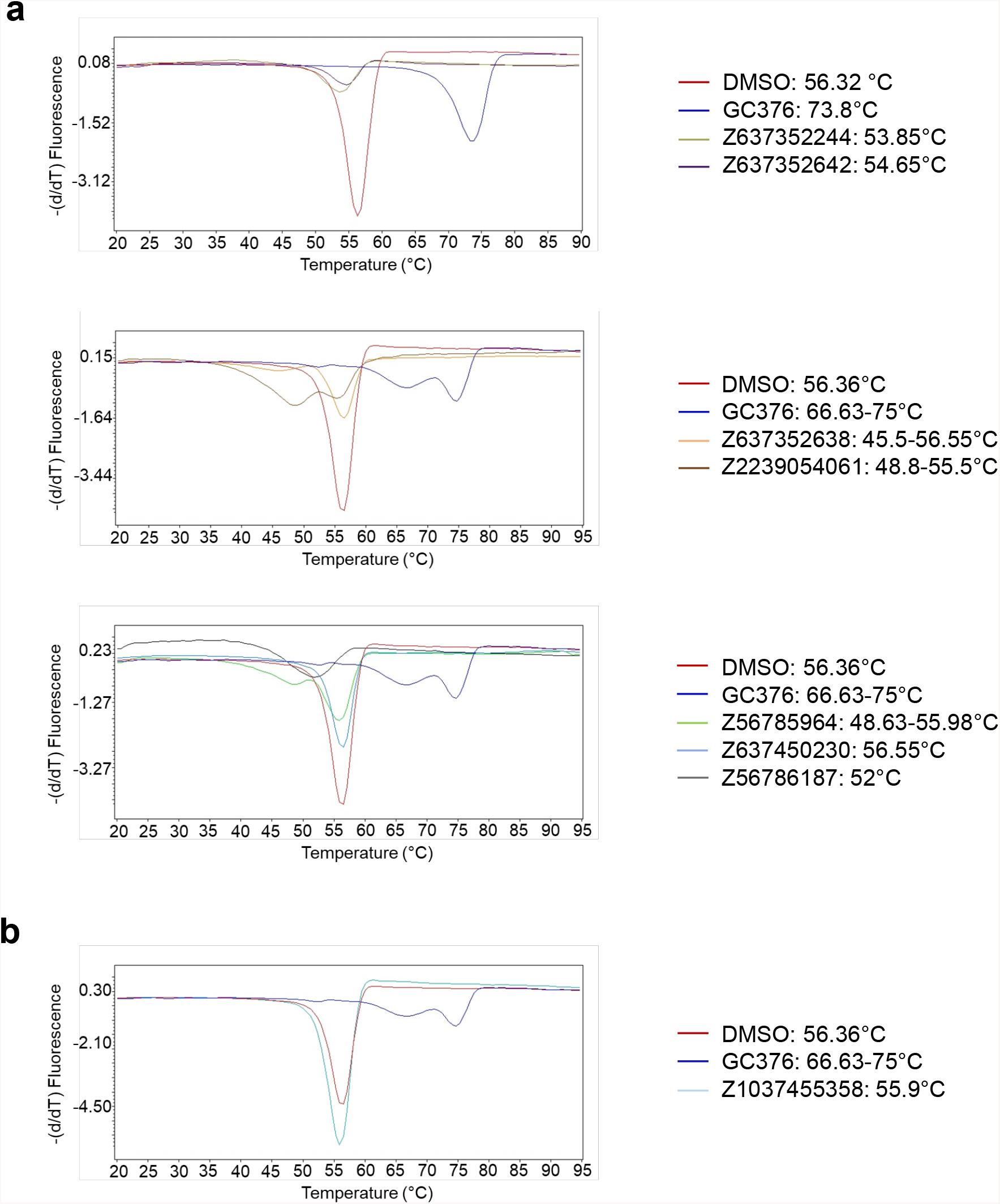
Effect of compounds containing nitrile or diamino-quinazoline groups on the melting temperature of M^pro^. Thermal shift assays were performed in the presence of DMSO or 20 μM M^pro^ inhibitors. The derivatives of the melting curves, shown in the graphs, were used to calculate the melting temperature of M^pro^. **a.** Compounds from the nitrile cluster. **b.** Diamino-quinazoline singleton Z1037455358.

The observed decrease in the melting temperature of M^pro^ by the above compounds was of concern, as it could indicate that these compounds were inhibiting the protease activity of M^pro^ by acting as unspecific denaturing agents. We reasoned that, if this was the case, then the compounds would lose inhibitory activity when the protease assay was performed in the presence of non-specific proteins that could serve as a sink for compound sequestration. Indeed, the parent nitrile-containing compound Z637352244, its analogues Z56786187 and Z637450230, and the Z1037455358 singleton all lost activity when the protease assay was performed in the presence of 1 μg cell lysate (Fig. 5). In contrast, the previously described inhibitor GC376 maintained activity in the presence of the lysate and, interestingly, so did compounds ZINC000373659060 and ZINC000636416501 (Fig. 5), which were identified by the second *in silico* screen (screen 2) [21].

**Fig. 5.**
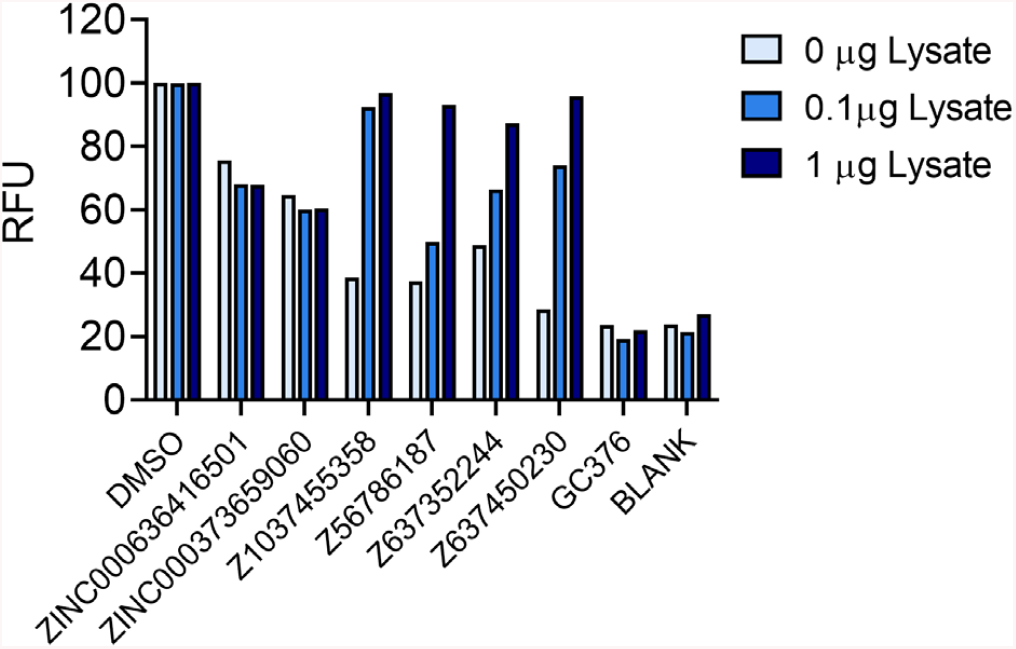
Effect of cell lysate on the activity of M^pro^ inhibiting compounds. The M^pro^ FRET protease assay was performed in the absence or presence of 0.1 or 1 μg cell lysate. The final concentration of the compounds was 40 μM.

### Characterization of a cluster of dihydro-quinolinone compounds

Compounds ZINC000373659060 and ZINC000636416501 are structurally related to each other and contain a dihydro-quinolinone core (Fig. 2b). Encouraged by the fact that the activity of these two compounds was not affected by the presence of cell lysate, we obtained 157 analogues (Table S7) and examined their ability to inhibit the protease activity of M^pro^. Three analogues were found to be significantly more potent than the parent compounds (Fig. 6a). Specifically, compounds Z228770960, Z393665558 and Z225602086 had IC_50_ values of 4.2, 5.9 and 7.4 μM, respectively (Fig. 6b). Importantly, all three analogues retained their inhibitory activity against M^pro^ in the presence of a cell lysate (Fig. 6c).

**Fig. 6.**
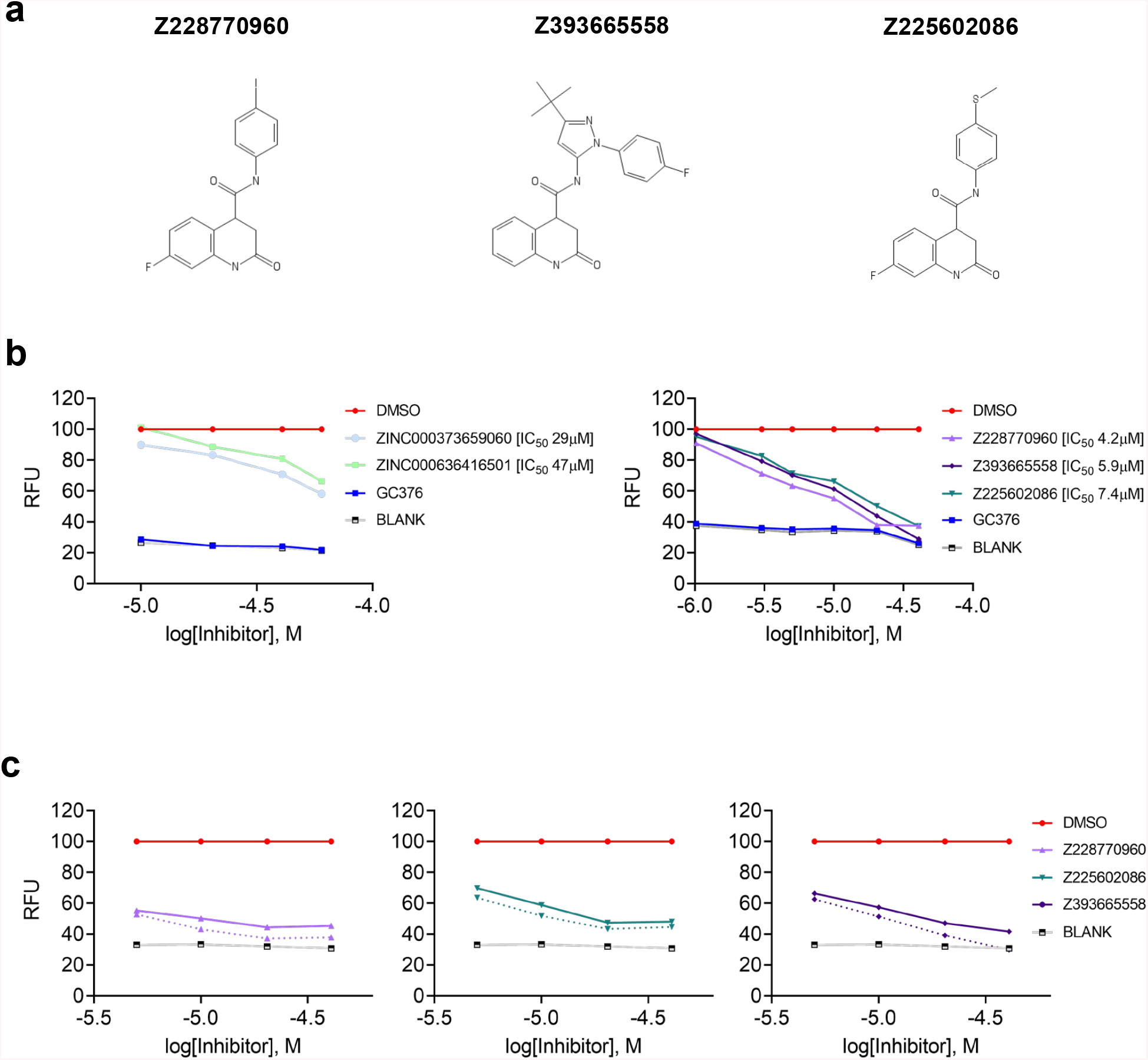
Dose-response curves of the dihydro-quinolinone compounds. **a**. Chemical structures of the most active dihydro-quinolinone compounds. **b.** Dose-response curves of the parent compounds ZINC000373659060 and ZINC000636416501 (10, 20, 40, 60 μM) and their analogues (1, 3, 5, 10, 20, 40 μM) in the M^pro^ protease assay. **c**. Dose-response curves of ZINC000373659060 analogues (5, 10, 20, 40 μM) in absence (continuous lines) or presence (dashed lines) of 0.1 μg protein lysate.

The parent ZINC000373659060 and ZINC000636416501 compounds and their three active analogues were then examined for their ability to modulate the melting temperature of M^pro^ in the thermal shift assay. The parent compounds did not affect the melting temperature of M^pro^ (Fig. 7a). However, the analogues increased the melting temperature of M^pro^ (Fig. 7b); the most active analogue, Z228770960, induced the greatest increase in melting temperature (1.2°C; Fig. 7b). These results are consistent with the hypothesis that the analogues inhibit M^pro^ by binding to its active site and stabilizing the protein, as predicted by the docking software that led to the identification of the parent compounds ZINC000373659060 and ZINC000636416501 (Fig. 8).

**Fig. 7.**
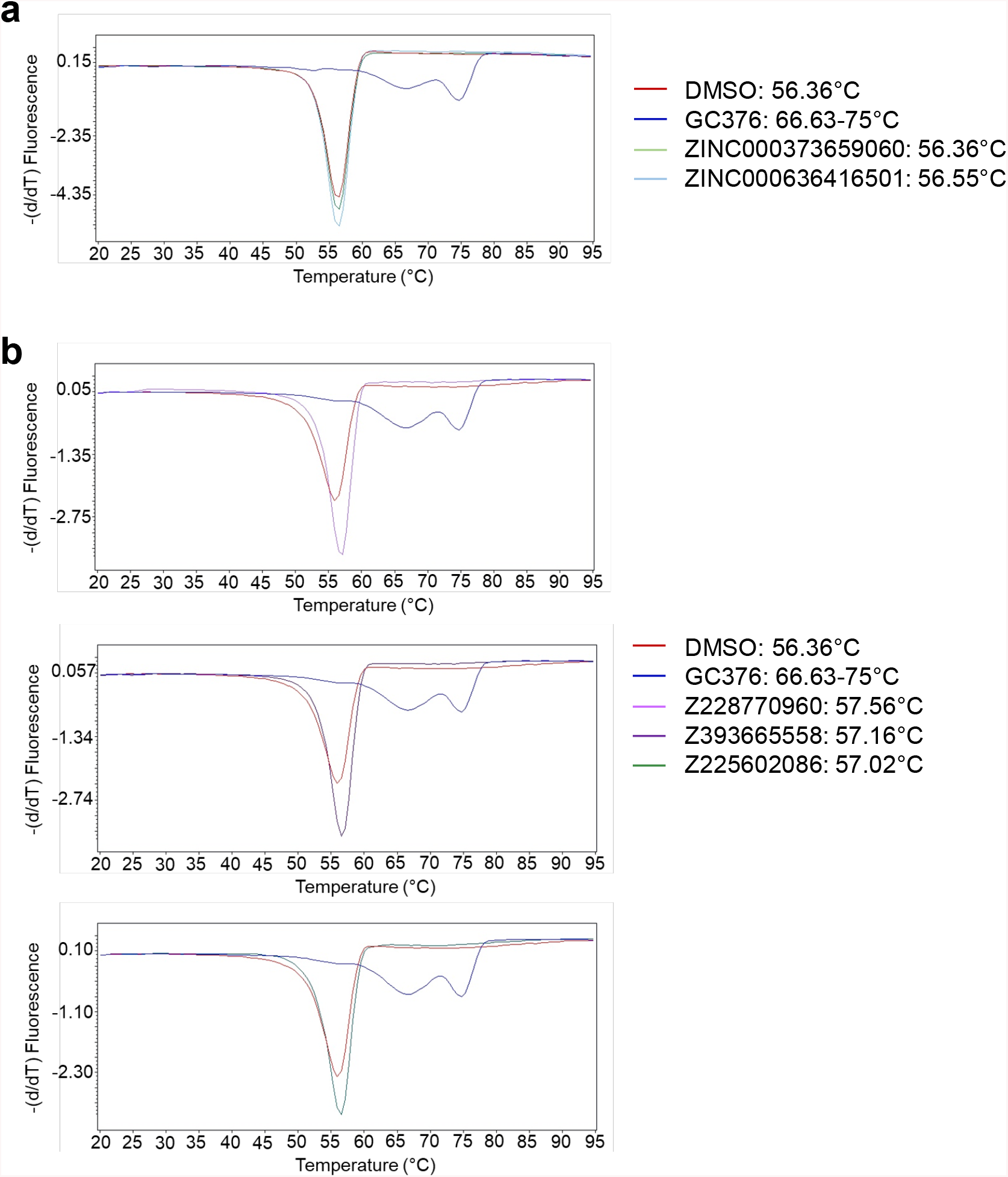
Effect of compounds from the dihydro-quinolinone cluster on the melting temperature of M^pro^. Thermal shift assays were performed in the presence of DMSO or 20 μM M^pro^ inhibitors. The derivatives of the melting curves, shown in the graphs, were used to calculate the melting temperature of M^pro^. **a.** Parent ZINC000373659060 and ZINC000636416501 compounds. **b.** Analogues.

**Fig. 8.**
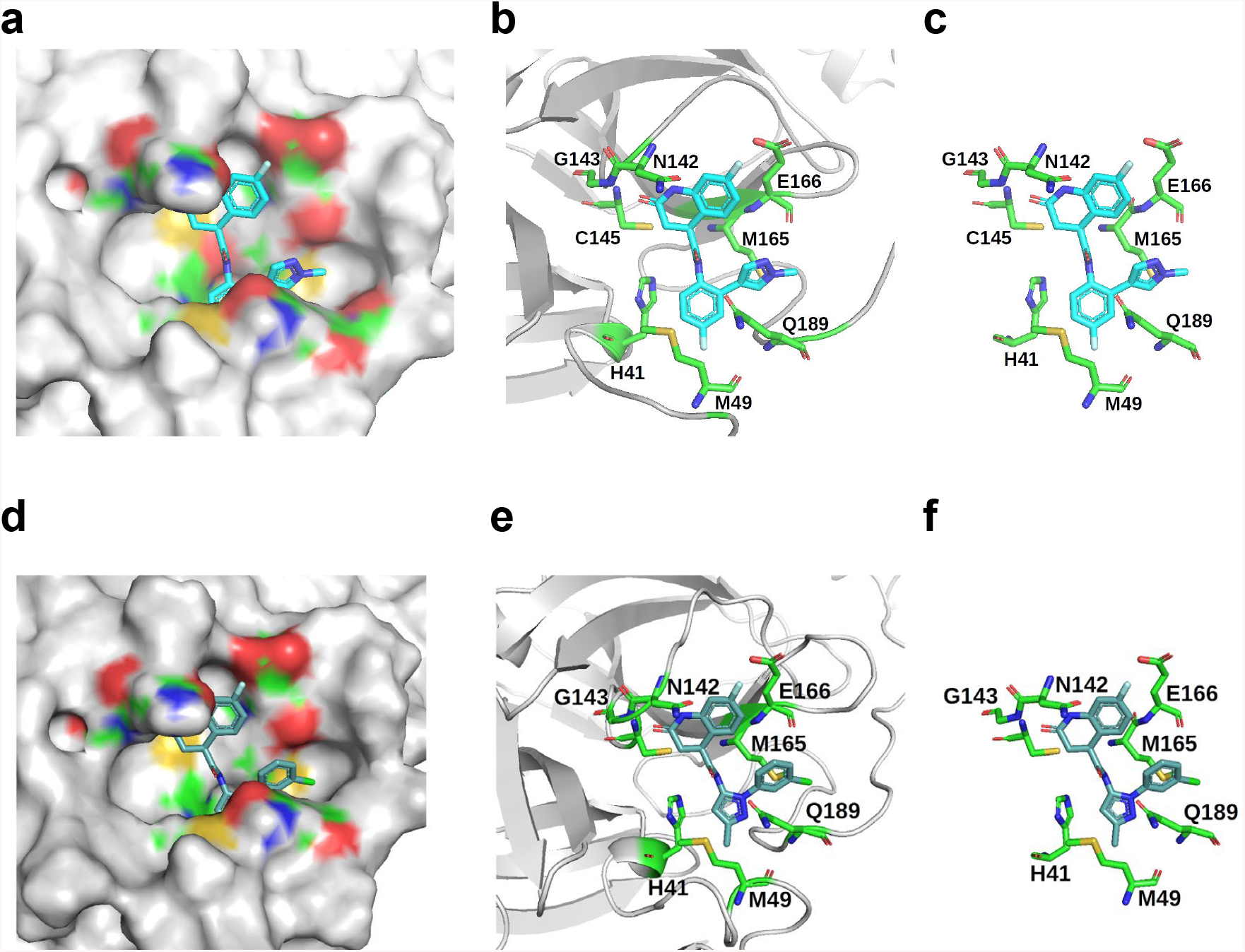
Docked conformations of parent dihydro-quinolinone compounds on M^pro^. **a-c**. Compound ZINC000373659060. **d-f.** Compound ZINC000636416501. **a,d.** Surface representation of SARS-CoV-2 M^pro^ protease with the predicted lowest energy docked pose of the compound shown as a stick model. **b,c,e,f.** Visualization of the residues at the active site of M^pro^ that potentially interact with the ligand. **b,e.** The secondary structure elements of M^pro^ are shown in cartoon form. All views are from the same orientation.

### Prospects

COVID-19 has had a significant impact on our society and this is likely to continue for several additional months. The development of vaccines against SARS-CoV-2 has proceeded with great speed, as compared to previously developed ones. Yet, despite their rapid development, the vaccines will become available only about one year after it became clear that COVID-19 was spreading as a pandemic and it will take several months to vaccinate a significant fraction of the population.

It is interesting to note that the active site of the M^pro^ protease of SARS-CoV-2 is identical to that of SARS-CoV-1 and very similar to those of the proteases of many other members of the coronavirus family. Indeed, the best SARS-CoV-2 M^pro^ inhibitors are actually compounds that were originally identified as inhibitors of SARS-CoV-1 M^pro^ [16,17,23]. However, these compounds were not developed to the point that they could be used to treat COVID-19 patients, because the SARS-CoV-1 epidemic was short-lived and the interest in developing agents that could treat SARS waned. In retrospect, it appears that this decision was a mistake.

The analogues that we have identified here are not potent enough to treat COVID-19 patients and their toxicity and pharmacokinetic profile is unknown. However, they have drug-like features and, therefore, the potential to be developed into drugs. In fact, these compounds might be the first class of compounds developed *de novo* against SARS-CoV-2 M^pro^ [23]. In addition, whereas the majority of M^pro^ inhibitors form covalent complexes with the catalytic cysteine, the compounds that we have identified are non-covalent inhibitors, lacking a highly reactive electrophile, which might make it easier to develop them for oral administration, a prerequisite for responding to a pandemic. For these reasons, we intend to proceed with the development of these compounds, not to impact the evolution of the current pandemic, but to better prepare ourselves for the next one.

## Supporting information

Supplementary_Tables_1-7__COVID-19_Mpro_02-12-2020

## Acknowledgements

The authors thank Anh-Tien Ton and Artem Cherkasov for providing the pdb coordinates of compounds ZINC000373659060 and ZINC000636416501 docked to M^pro^ [21]. Funding for this project was provided by the Fondation Aclon and by overhead funds from the ERC Project ONIDDAC (Principal Investigator: T.D.H.).

## BIBLIOGRAPHY

[1] N. Zhu, D. Zhang, W. Wang, X. Li, B. Yang, J. Song, X. Zhao, B. Huang, W. Shi, R. Lu, P. Niu, F. Zhan, X. Ma, D. Wang, W. Xu, G. Wu, G. F. Gao, and W. Tan for the China Novel Coronavirus Investigating and Research Team, N Engl J Med 2020, 382, 727–733

[2] A. E. Gorbalenya, S. C. Baker, R. S. Baric, R. J. de Groot, C. Drosten, A. A. Gulyaeva, B. L. Haagmans, C. Lauber, A. M. Leontovich, B. W. Neuman, D. Penzar, S. Perlman, L. L. M. Poon, D. V. Samborskiy, I. A. Sidorov, I. Sola & J. Ziebuhr, Nat Microbiol 2020, 5, 536–544

[3] W. Dai, B. Zhang, X. Jiang, H. Su, J. Li, Y. Zhao, X. Xie, Z. Jin, J. Peng, F. Liu, C. Li, Y. Li, F. Bai, H. Wang, X. Cheng, X. Cen, S. Hu, X. Yang, J. Wang, X. Liu, G. Xiao, H. Jiang, Z. Rao, L. Zhang, Y. Xu, H. Yang, H. Liu, Science 2020, 368 (6497, 1331–1335

[4] L. Zhang, D. Lin, X. Sun, U. Curth, C. Drosten, L. Sauerhering, S. Becker, K. Rox, R. Hilgenfeld, Science 2020, 368 (6489, 409–412

[5] Z. Jin, X. Du, Y. Xu, Y. Deng, M. Liu, Y. Zhao, B. Zhang, X. Li, L. Zhang, C. Peng, Y. Duan, J. Yu, L. Wang, K. Yang, F. Liu, R. Jiang, X. Yang, T. You, X. Liu, X. Yang, F. Bai, H. Liu, X. Liu, LW. Guddat, W. Xu, G. Xiao, C. Qin, Z. Shi, H. Jiang, Z. Rao, H. Yang, Nature 2020, 582 (7811, 289–293

[6] H. Su, S. Yao, W. Zhao, M. Li, J. Liu, W. Shang, H. Xie, C. Ke, H. Hu, M. Gao, K. Yu, H. Liu, J. Shen, W. Tang, L. Zhang, G. Xiao, L. Ni, D. Wang, J. Zuo, H. Jiang, F. Bai, Y. Wu, Y. Ye & Y. Xu, Acta Pharmacol Sin 2020, 41, 1167–1177

[7] Z. Jin, Y. Zhao, Y. Sun, B. Zhang, H. Wang, Y. Wu, Y. Zhu, C. Zhu, T. Hu, X. Du, Y. Duan, J. Yu, X. Yang, X. Yang, K. Yang, X. Liu, L. W. Guddat, G. Xiao, L. Zhang, H. Yang & Z. Rao, Nat Struct Mol Biol 2020, 27, 529–532

[8] H. Su, S. Yao, W. Zhao, M. Li, J. Liu, W. Shang, H. Xie, C. Ke, H. Hu, M. Gao, K. Yu, H. Liu, J. Shen, W. Tang, L. Zhang, G. Xiao, L. Ni, D. Wang, J. Zuo, H. Jiang, F. Bai, Y. Wu, Y. Ye & Y. Xu, Acta Pharmacol Sin 2020, 41, 1167–1177

[9] C.D. Owen, P. Lukacik, C.M. Strain-Damerell, A. Douangamath, A. J. Powell, D. Fearon, J. Brandao-Neto, A. D. Crawshaw, D. Aragao, M. Williams, R. Flaig, D. R. Hall, K. E. McAuley, M. Mazzorana, D. I. Stuart, F. von Delft, M. A. Walsh, PDB 2020, ID: 6YD7 DOI: 10.2210/pdb6yb7/pdb

[10] Y. Kim, S. Lovell, K. Tiew, S. R. Mandadapu, K. R. Alliston, K. P. Battaile, W. C. Groutas, K. Chang, Journal of virology 2012, 86 (21, 11754–11762

[11] K. Tiew, G. He, S. Aravapalli, S. R. Mandadapu, M. R. Gunnam, K. R. Alliston, G. H. Lushington, Y. Kim, K. Chang, W. C. Groutas, Bioorg Med Chem Lett 2011, 21, 5315–5319

[12] S. R. Mandadapua, P. M. Weerawarna, M. R. Gunnam, K. R. Alliston, G. H. Lushington, Y. Kim, K. Chang, and W. C. Groutas, Bioorg Med Chem Lett 2012, 22, 4820–4826

[13] Y. Kim, V. Shivanna, S. Narayanan, A. M. Prior, S. Weerasekara, D. H. Hua, A. C. G. Kankanamalage, W. C. Groutas, K. Chang, Journal of virology 2015, 89, 4942–4950

[14] W. Vuong, M. B. Khan, C. Fischer, E. Arutyunova, T. Lamer, J. Shields, H. A. Saffran, R. T. McKay, M. J. van Belkum, M. A. Joyce, H. S. Young, D. L. Tyrrell, J. C. Vederas & M. J. Lemieux, Nat Commun 2020, 11, 4282

[15] C. Ma, M. D. Sacco, B. Hurst, J. A. Townsend, Y. Hu, T. Szeto, X. Zhang, B. Tarbet, M. T. Marty, Y. Chen & J. Wang, Cell Res 2020, 30, 678–692

[16] R. L. Hoffman, R. S. Kania, M. A. Brothers, J. F. Davies, R. A. Ferre, K. S. Gajiwala, M. He, R. J. Hogan, K. Kozminski, L. Y. Li, J. W. Lockner, J. Lou, M. T. Marra, L. J. Mitchell Jr., B. W. Murray, J. A. Nieman, S. Noell, S. P. Planken, T. Rowe, K. Ryan, G. J. Smith III, J. E. Solowiej, C. M. Steppan, and B. Taggart, J Med Chem 2020, 63 (21, 12725–12747

[17] B. Boras, R. M. Jones, B. J. Anson, D. Arenson, L. Aschenbrenner, M. A. Bakowski, N. Beutler, J. Binder, E. Chen, H. Eng, J. Hammond, R. Hoffman, E. P. Kadar, R. Kania, E. Kimoto, M. G. Kirkpatrick, L. Lanyon, E. K. Lendy, J. R. Lillis, S. A. Luthra, C. Ma, S. Noell, R. S. Obach, M. N. O’ Brien, R. O’Connor, K. Ogilvie, D. Owen, M. Pettersson, M. R. Reese, T. F. Rogers, M. I. Rossulek, J. G. Sathish, C. Steppan, M. Ticehurst, L. W. Updyke, Y. Zhu, J. Wang, A. K. Chatterjee, A. D. Mesecar, A. S. Anderson, C. Allerton, bioRxiv 2020, 2020.09.12.293498

[18] A. Douangamath, D. Fearon, P. Gehrtz, T. Krojer, P. Lukacik, C. D. Owen, E. Resnick, C. Strain-Damerell, A. Aimon, P. Ábrányi-Balogh, J. Brandão-Neto, A. Carbery, G. Davison, A. Dias, T. D. Downes, L. Dunnett, M. Fairhead, J. D. Firth, S. P. Jones, A. Keeley, G. M. Keserü, H. F. Klein, M. P. Martin, M. E. M. Noble, P. O’Brien, A. Powell, R. N. Reddi, R. Skyner, M. Snee, M. J. Waring, C. Wild, N. London, F. von Delft & M. A. Walsh, Nat Commun 2020, 11, 5047

[19] D. Zaidman, P. Gehrtz, M. Filep, D. Fearon, J. Prilusky, S. Duberstein, G. Cohen, D. Owen, E. Resnick, C. Strain-Damerell, P. Lukacik, H. Barr, M.A. Walsh, F. von Delft, N. London, Covid-Moonshot Consortium, bioRxiv 2020, doi: 10.1101/2020.09.21.299776.

[20] C. Gorgulla, K.M. Padmanabha Das, K.E. Leigh, M. Cespugli, P.D. Fischer, Z. Wang, G. Tesseyre, S. Pandita, A. Shnapir, A. Calderaio, C. Hutcheson, M. Gechev, A. Rose, N. Lewis, E. Yaffe, R. Luxenburg, H.D. Herce, V. Durmaz, T.D. Halazonetis, K. Fackeldey, J.J. Patten, A. Chuprina, I. Dziuba, A. Plekhova, Y. Moroz, D. Radchenko, O. Tarkhanova, I. Yavnyuk, C.C. Gruber, R. Yust, D. Payne, A.M. Näär, M.N. Namchuk, R.A. Davey, G. Wagner, J. Kinney, H. Arthanari, ChemRxiv 2020, doi: 10.26434/chemrxiv.12682316.v1.

[21] A. Ton, F. Gentile, M. Hsing, F. Ban, A. Cherkasov, Molecular Informatics 2020, 39 (8), e2000028

[22] M.C. Lo, A. Aulabaugh, G. Jin, R. Cowling, J. Bard, M. Malamas, G. Ellestad, Anal Biochem 2004, 332, 153–159.

[23] Y. Liu, C. Liang, L. Xin, X. Ren, L. Tia, X. Ju, H. Li, Y. Wang, Q. Zhao, H. Liu, W. Cao, X. Xie, D. Zhang, Y. Wang, Yanlin Jian, Eur J Med Chem 2020, 206, 112711

